# Inhibiting fibronectin assembly in the breast tumor microenvironment increases cell death and improves response to doxorubicin

**DOI:** 10.1101/2025.02.12.637963

**Authors:** Metti K. Gari, Hye Jin Lee, David R. Inman, Brian M. Burkel, Margaret A. Highland, Glen S. Kwon, Nikesh Gupta, Suzanne M. Ponik

## Abstract

**Purpose:** Effective therapies for solid tumors, including breast cancers, are hindered by several roadblocks that can be largely attributed to the fibrotic extracellular matrix (ECM). Fibronectin (FN) is a highly upregulated ECM component in the fibrotic tumor stroma and is associated with poor patient prognosis. This study aimed to investigate the therapeutic potential of an anti-fibrotic peptide that specifically targets FN and blocks the fibrillar assembly of FN.

**Methods:** To target FN, we used PEGylated Functional Upstream Domain (PEG-FUD), which binds to the 70 kDa N-terminal region of FN with high affinity, localizes to mammary tumors, and potently inhibits FN assembly in vitro and in vivo. Here, we used the 4T1 tumor model to investigate the efficacy and mechanisms of PEG-FUD to inhibit tumor growth.

**Results:** Our data demonstrates that PEG-FUD monotherapy reduces tumor growth without systemic toxicity. Analysis of the tumor microenvironment revealed that PEG-FUD effectively inhibited FN matrix assembly within tumors and reduced adhesion-mediated signaling through α5 integrin and FAK leading to enhanced tumor cell death. Notably, signaling through FAK has been associated with resistance mechanisms to doxorubicin (DOX). Therefore, we tested the combination of PEG-FUD and Dox, which significantly reduced tumor growth by 60% compared to vehicle control and 30% compared to Dox monotherapy.

**Conclusions:** Our findings demonstrate that PEG-FUD significantly modifies the peritumoral ECM of breast cancer, leading to increased tumor cell death, and potentiates the efficacy of conventional breast cancer therapy.

## Introduction

Therapies for solid tumors, including breast cancer, are hindered by several roadblocks that can be largely attributed to the fibrotic ECM in the microenvironment of these tumors. The fibrotic ECM in the breast tumor microenvironment (TME) is well-known for promoting tumor aggressiveness, metastatic progression, and mechanisms of therapy resistance(1–3). During tumor progression, the deposition of ECM proteins such as collagen and FN is significantly increased, which results in tumor fibrosis. Importantly, the excess deposition of ECM in the breast TME along with alterations in ECM architecture are associated with poor breast cancer patient outcome(4–9). Based on the abundance of ECM in solid tumors and the importance of the fibrotic ECM in breast cancer progression, multiple efforts have been made to therapeutically target the ECM. To date, ECM targeting strategies have focused on interfering with ECM-modulating enzymes or signaling pathways involved in the deposition, degradation, and remodeling of ECM components (10–13). Despite the promising pre-clinical success of existing ECM-targeting drugs, most fail to transition to the clinic due to toxicity and off-target effects(14–17). Hence, there is a need to develop new therapies that specifically target the ECM and are potent enough to achieve efficacy while maintaining safety.

In this study, we focus on directly targeting FN, one of the major ECM proteins identified in the fibrotic tumor matrix(18–20). FN is a multifunctional glycoprotein that exists in two major forms, plasma FN secreted from the liver and cellular FN that is secreted locally by various cell types (21). Despite the source, FN is secreted as a soluble dimer in a compact inactive state, where it can be incorporated into an insoluble fibrillar matrix upon binding to cell surface receptors, including the integrin receptor α5β1(22). The N-terminal 70-kDa region of FN is required for the assembly of FN into an insoluble fibrillar matrix(23). Insoluble or assembled FN matrix in the TME plays a crucial role in scaffolding other ECM proteins, stabilizing the ECM, and providing adhesion sites to initiate tumor cell signaling cascades that enhance proliferation and viability(24–26). To specifically target FN and block the fibrillar assembly of FN, we chose to use the small 6-kDa bacterial-derived peptide called Functional Upstream Domain (FUD/pUR4) discovered by Tomasini-Johansson et al(27). FUD binds to the N-terminal 70-kDa region of both sources of FN (cellular and plasma) with nanomolar binding affinity, consequently, blocking the extension of FN and assembly into fibrils(27). Importantly, FUD is a potent inhibitor of FN assembly both *in vitro* and *in vivo*(28–30). Several laboratories have demonstrated the potential of FUD as a therapeutic agent in multiple fibrotic diseases, including breast cancer(31). However, the peptide is rapidly cleared through the renal system, which limits the *in vivo* bioavailability and results in daily required FUD administration at a relatively high dose of ∼25 mg/kg(32). To address the issue of rapid renal clearance, we PEGylated-FUD (PEG-FUD) and demonstrated a significant increase in peptide bioavailability and tumor retention compared to non-PEGylated FUD, while maintaining the ability to inhibit FN assembly(33–36).

Here, we investigated the anti-cancer efficacy of PEG-FUD in the TNBC 4T1 mouse mammary carcinoma model. We hypothesized that modulating FN assembly into a fibrillar matrix with PEG- FUD would disrupt cell-ECM interactions thereby inhibiting tumor growth, blocking mechanisms of drug resistance, and improving the efficacy of chemotherapy. Our study revealed that treatment with PEG-FUD significantly decreased 4T1 tumor growth. By characterizing the alterations in the TME in response to PEG-FUD, we identified a significant inhibition of FN matrix assembly, reduced levels of cell adhesion proteins (including α5 integrin and FAK), and an increase in caspase-3 mediated cell death. Further investigation into the potential of PEG-FUD to improve chemotherapy response revealed that co-treatment with PEG-FUD significantly enhanced the anti-tumor efficacy of Dox without any additional systemic toxicity.

## Materials and Methods

### Preparation of Functional Upstream Domain (FUD) peptide and its mutated counterparts

FUD and the FUD control (<u>m</u>utated FUD (mFUD) and <u>F</u>ully <u>m</u>utated FUD (FmFUD)) constructs were a kind gift from Dr. Donna Peters at University of Wisconsin-Madison. The mFUD and FmFUD control peptides have reduced binding affinity for FN due to point mutations in the FN binding regions. All peptide sequences were cloned into the pET-ELMER vector and recombinantly expressed in BL21 *Escherichia coli* (DE3) as His-tagged peptides as previously described by our group(33–36). Briefly, 1 mM of isopropyl β-D-1thiogalactopyranoside (IPTG) was added to the obtained 1L culture to induce expression of FUD/mFUD/FmFUD. The resulting bacterial pellet was suspended in the lysis buffer (pH 8.0) [Urea 8M, NaH_2_PO_4_ 100 mM, Tris 10 mM, Imidazole 5 mM] and centrifuged to obtain clear supernatant. Ni-NTA agarose (Qiagen) beads were equilibrated with the lysis buffer and 1 mL of Ni-NTA agarose resin was added for every 4 mL of the above obtained lysate and incubated overnight at 4°C with Ni-NTA agarose beads. The next day, the lysate was removed and the Ni-NTA agarose was washed with a washing buffer (pH 8.0) [Urea 8M, NaH_2_PO_4_ 100 mM, Tris 10 mM, Imidazole 15 mM] and equilibrated with Thrombin cleavage buffer (pH 8.4) [CaCl_2_ 2.5 mM, NaCl 150 mM, Tris 20 mM]. Elution of peptides and removal of the His tag was achieved by using a thrombin cleavage site between the His-tag and the corresponding peptide (FUD/mFUD/FmFUD). FUD bound to the Ni-NTA agarose was incubated with ∼36 units of Bovine α-Thrombin for every 15 mL of sample. The eluted sample was collected and further purified via fast protein liquid chromatography (FPLC) using HiTrap Q HP column from Cytiva.

### PEGylation of FPLC purified peptides

PEGylated conjugates were synthesized using reductive amination chemistry. FUD, mFUD or FmFUD peptide(s) were N-terminally selectively conjugated with 20-kDa methoxy-PEG- propionaldehyde using the previously published protocol by our group(33). Briefly, the FPLC purified FUD/mFUD/FmFUD peptide was mixed with PEG in a sodium acetate buffer solution (50 mM, pH 5.5) in the presence of NaBH_3_CN and the reaction was carried out for ∼18 hr in the fuming cabinet under cold conditions (4°C). After the reaction time was complete, the mixture was dialyzed two times against NaOAc buffer to remove excess reducing agent. Next, the buffer was switched to 20mM Tris and left overnight before FPLC purification via ion-exchange chromatography using 5mL HiTrap Q HP anion exchange column. The concentration of FUD/mFUD/FmFUD was obtained by measuring the absorbance at 280 nm using corresponding ε values [0.496 (FUD), 0.742 (mFUD) and 0.250 (FmFUD)]. Patent number for PEG-FUD: US10828372B2.

### Enzyme-linked competition binding assay

The enzyme-linked competition binding assay was carried out as previously reported(37,38), with a small modification. Briefly, a 96-well high-binding culture plate was coated overnight with human plasma FN under cold conditions (4°C) at 10 μg/mL of concentration. The biotinylated FUD (b-FUD) was prepared as per the manufacturer’s protocol using NHS-biotin-ester (Pierce). The above FN-coated plate was blocked for 1 hr using 5% BSA prepared in tris-buffered saline containing 0.05% Tween 20 (TBS-T). 0.5 nM of b-FUD was added to the plate concurrently with various concentration (2000, 1000, 500, 250, 125, 62.5, 0 nM) of unlabeled PEG-FmFUD or PEG- FUD in 0.1% BSA in TBS-T. After a 2 hr incubation at RT, followed by three TBS-T washes, alkaline phosphatase-conjugated streptavidin (Jackson Immunoresearch) was added at 1:20,000 dilution factor and incubated for 1 hr at RT. After washing with TBS-T again, 100 uL of 1-step p-Nitrophenyl Phosphate was added as a substrate to each treated well, and the reaction was stopped by the addition of 2N sodium hydroxide. Absorbance was measured at 405 nm using a microplate reader.

### Cell monolayer culture

4T1 breast cancer cells were obtained from American Type Culture Collection (VA). 4T1 cells were cultured in Rosewell Park Memorial Institute Medium-1640 (RPMI) (Corning, NY, USA) containing 10% fetal bovine serum (FBS). The human breast cancer-associated fibroblasts (hCAFs) cells were a gift from Charlotte Kupperwaser’s laboratory at Tufts University. The hCAFs were cultured in Dulbecco’s Modified Eagle Medium (DMEM, Gibco) containing 10% FBS and 1% streptomycin/penicillin. The bone-marrow-derived fibroblast cells isolated from the femurs of BALB/c mice were cultured in DMEM containing 20% FBS and 1% streptomycin/penicillin and activated by 4T1-conditioned media to generate murine cancer-associated fibroblasts (mCAFs) as described previously (39). All cell cultures were kept at 37°C in the humidified incubator with 5% CO_2_. For tumor cell inoculations, cells were washed, harvested and counted manually using a hemocytometer.

### Cell monolayer ECM staining

To confirm ECM changes in fibroblast culture after PEG-FUD incubation, 25,000 hCAFs or mCAFs in corresponding media containing 50 μg/ml of ascorbic acid were seeded onto a 35 mm glass bottom dish (VWR). The next day, 250 nM of PEG-FUD peptides in phosphate-buffered saline (PBS) were incubated with hCAFs or 500 nM of PEG-FUD was incubated with mCAFs for 48 hr. PBS was used as a vehicle in control groups. After the peptide incubation, cells were washed with Hanks’ Balanced Salt Solution and fixed with 10% formalin for 10 min at RT. Following fixation, 2% BSA in PBS was used for blocking for 1 hr at RT. hCAFs were incubated with a A488-labeled anti-FN primary antibody (eBioscience, No. 53-9869-82, 1:100) in 1% BSA/PBS at 4°C overnight. The next day, cells were washed to remove unbound primary antibody. mCAFs, were immunolabeled using a rabbit anti-mouse FN primary antibody (1:500) (34) in 1% BSA/PBS at 4°C overnight and an A488-labeled goat anti-rabbit (Life technologies, A11034, 1:500) secondary antibody in 1% BSA in PBS for 1 hr at RT. After washing, nuclei were stained with DAPI (D9542, MilliporeSigma). Confocal microscopy (Nikon AⅠRS with 20x objective lens) was used to visualize the assembled FN by the fibroblasts, and images were processed with the vendor software (NIS-Elements).

### Animal Tumor model

9-week-old female BALB/c mice were purchased from Jackson Labs and housed in the Wisconsin Institutes for Medical Research animal facilities at the University of Wisconsin-Madison. All experiments were conducted in compliance with the University of Wisconsin Institutional Animal Care and Use Committee (IACUC). 4T1 tumors were established by orthotopically injecting 2×10^5^ 4T1 cells in 50 µl DPBS into the fourth mammary fat pad of syngeneic, immunocompetent BALB/c mice. To minimize tumor abrasion, mice were housed on soft bedding. To assess FN protein levels, major organs such as the mammary glands, lungs, liver, spleen, and kidney were harvested from 4T1 tumor bearing and non-tumor bearing control mice 4 weeks after 4T1 cell inoculation. For PEG-FUD monotherapy, after orthotopic tumor injection, all mice were randomly assigned to an experimental group. After 1 week of tumor growth, when tumor size reached approximately 50 mm^3^, vehicle control (Ctrl_PBS) or peptides (PEG-FmFUD, PEG-mFUD and PEG-FUD) were administered subcutaneously at the scruff of the neck every 48 hr at a dose of 12.5mg/kg (outlined in Fig 3). For combination therapy (PEG-FUD + Dox), 4 doses of PEG-FUD or control peptides were administered every 48 hrs, then Dox treatment was initiated, which consisted of intraperitoneal administration every 72 hr at 5 mg/kg along with continued peptide dosing every 48 hrs (outlined in Fig 7A). During the treatment course, tumor growth was measured manually every 48 hr using a caliper, starting at the first treatment. Tumor volume was calculated using the formula V = ½ (Length × Width^2^)(40). The maximal tumor volume approved by the IACUC at the University of Wisconsin-Madison is 1.5 cm^3^. In compliance with these guidelines, all animals in this study were euthanized at or before tumors reached this volume (Supplemental Table 1A-B). At the end of the treatment, blood was collected, mice were euthanized, and tissues were harvested for subsequent analysis.

### Toxicity evaluation

Body weight was monitored every 48 hr during the treatment course. At the experimental end point, blood was collected via cardiac puncture to perform a complete blood count (CBC) analysis using the VETSCAN® HM5 Hematology Analyzer. During harvest, the spleen was collected and weighed. Additionally, kidneys and liver were harvested and fixed in 10% buffered formalin for 48 hr at 4 °C. Fixed tissues were paraffin embedded and 5 μm sections were stained with hematoxylin and eosin (H&E) by routine methods. Stained slide sections were captured at 40x by Aperio Digital Pathology Slide Scanner system and the digitized slides were evaluated by a board certified veterinary anatomic pathologist using Aperio ImageScope [v12.4.6.5003].

### Immunofluorescence staining

4T1 tumors were fixed for 48 hr at 4 °C in 10% buffered formalin. Fixed tissues were paraffin-embedded and sectioned into 5 μm sections for immunofluorescence analysis. Briefly, tissue sections were deparaffinized in a humidity chamber with xylene and rehydrated in graded ethanol washes. Next tissue sections were subjected to heat-induced antigen retrieval in citrate buffer pH 6.0. Slides were blocked with 10% BSA in TBS for 1 hr then incubated for 1 hr at RT with primary antibodies, FN (ab23750, Abcam, 1:250) and Antigen Kiel 67- Ki67 (ab15580, Abcam, 1:100), followed by washing in TBS-T. Next, samples were incubated with secondary antibody (ab7090, Abcam, 1:500) for 10 min at RT. For amplified fluorescence detection, sections were incubated with TSA substrates (PerkinElmer) for 10 min at RT, followed by nuclear counterstaining with DAPI (D9542, MilliporeSigma). Slides were coverslipped using ProLong Gold (no. P36930; Thermofisher). Immunofluorescent labeled slides were imaged using a Leica Thunder imaging system with 10x magnification. Quantification of >10 fields of view was performed in Image J for each staining.

### Picro-sirius red (PSR) staining

Formalin fixed paraffin embedded (FFPE) 4T1 tumor tissues were sectioned at 5 μm for PSR staining. Briefly, tissue sections were deparaffinized with xylene and underwent sequential washes of graded ethanol as described above. Next slides were stained with PSR (Sigma-Aldrich; "Direct Red 80"; Catalog # 36-554-8) for 1 hr. Slides were then washed with acidified water, dehydrated in 100 % ethanol, and cleared in xylene. Slides were coverslipped using Richard-Allan Scientific Mounting Medium (REF 4112, Thermo Scientific). PSR stained slides were imaged at an excitation wavelength of 575 nm using Leica Thunder imaging system with a magnification of 20x. Quantification of >10 fields of view were performed in Image J.

### TUNEL staining

FFPE 4T1 tumor sections were deparaffinized and rehydrated as described above. Sections were then stained with a 1:10 dilution of TUNEL enzyme in TUNEL buffer (Roche In Situ Cell Detection Kit w/Fluorescein. Catalog # 11684795910) for 1 hr at 37°C, washed with PBST, and coverslipped with ProLong Gold mounting media (no. P36930; Thermofisher). TUNEL stained slides were imaged at an excitation wavelength of 475 nm using Leica Thunder imaging system with a magnification of 20x.

### Western Blotting

Tissues were fractionated into soluble and insoluble lysates similar to our prior study(34). Briefly, tissues were cryopulverized using a mortar and pestle in the presence of liquid nitrogen. The homogenized samples were lysed in RIPA buffer containing 1% deoxycholate (DOC) at 0.1 g tissue per ml buffer and spun at 4°C. The supernatant was collected, and the pellets were solubilized in buffer containing 4 M urea, 4% SDS and 1 mM DTT. Resuspended pellets were vortexed, sonicated, and heated to 95°C for 5 min. The DOC-soluble supernatant contains nuclear, cytoplasmic, and membrane fractions, while mainly ECM proteins constitute the DOC-insoluble pellet. Protein concentrations were measured using the DC Protein Assay kit (Bio-Rad) and BSA standards, diluted in the corresponding buffers. Proteins in each fraction were separated by SDS– polyacrylamide gel electrophoresis and transferred to a polyvinylidene difluoride membrane. To ensure even protein loading and transfer efficiency, total protein (TP) staining (Fast Green FCF, MilliporeSigma) was performed. Next, membrane was blocked with 5% non-fat dry milk solution in TBS-T, and incubated overnight at 4°C with the following primary antibodies: FN (ab23750, Abcam, 1:250, 2hr); COL1A1 (NBP1-30054, Novus Biologicals, 1:500); tenascin-C (ab108930, Abcam, 1:500); periostin ( ab14041, Abcam, 1:500) ; proliferating cell nuclear antigen – PCNA (#13110, Cell Signaling, 1:500); Caspase-3 (#9662, Cell Signaling, 1:500); integrin α5 (10569-1-AP, proteintech, 1:1000); integrin β1 (#34971, Cell Signaling, 1:1000); FAK ( #05-537, MilliporeSigma, 1:1000). Proteins were detected using either a horseradish peroxidase–conjugated secondary antibodies (Jackson ImmunoResearch) followed by chemiluminescence substrate (LI-COR BioScience) or fluorescent–conjugated secondary antibodies (IRDye infrared dyes from LI-COR Biosciences). Blots were imaged using the Odyssey Fc Imager and band intensities were analyzed using Image studio. For detection of plasma FN, blood was isolated from mice via cardiac puncture and spun for 10 min at 2,000 x-g to remove cells. Next, the plasma layer was collected and spun for 15 min at 2,000 x-g to deplete platelets in the plasma. The supernatant was collected and put in 2x Laemmli followed by standard procedures of Western blotting as described above.

### Statistics

GraphPad Prism was used to generate graphs and perform statistical analysis. For comparison between two groups, an unpaired t-test was performed. For comparison between multiple groups, a one-way analysis of variance (ANOVA) with Tukey’s Multiple Comparison Test (TMCT) was performed, with a corrected P value ≤0.05 considered significant. Data is reported as mean ± s.e.m.

## Results

### PEG-FUD inhibits mammary CAF-mediated FN assembly *in vitro*

Cancer-associated fibroblasts (CAF) are one of the most abundant stromal cells in the TME and the primary cell type responsible for the deposition of ECM proteins, including FN. CAFs not only secret FN but also assemble FN(41). We previously showed that Cy5-labeled PEG-FUD can bind to exogenous or endogenous FN assembled by CAFs(36). Prior studies have shown that PEG-FUD inhibits FN assembly by human foreskin fibroblasts(33,42), however, PEG-FUD has yet to be demonstrated as an inhibitor of FN assembly by mammary CAFs. To confirm PEG-FUD inhibits mammary CAF-mediated FN matrix assembly, we induced ECM matrix assembly in hCAFs or activated mCAFs and treated cells with control (Ctrl_PBS) or PEG-FUD (250 or 500 nM, respectively) for 48 hrs. At experimental endpoint, the cultures were fixed and the fibrillar structure of FN was visualized by immunofluorescence staining. PEG-FUD significantly decreased the area of cellular FN assembled by mCAFs (Fig 1A-B). Additionally, we identified a similar decrease in hCAF-mediated fibrillar FN assembly in the presence of PEG-FUD compared to the control, PBS-treated condition (Fig 1C-D). These results provide evidence that PEG-FUD broadly inhibits *in vitro* FN assembly by fibroblasts, including FN assembled by mammary CAFs *in vitro*.

**Figure 1:**
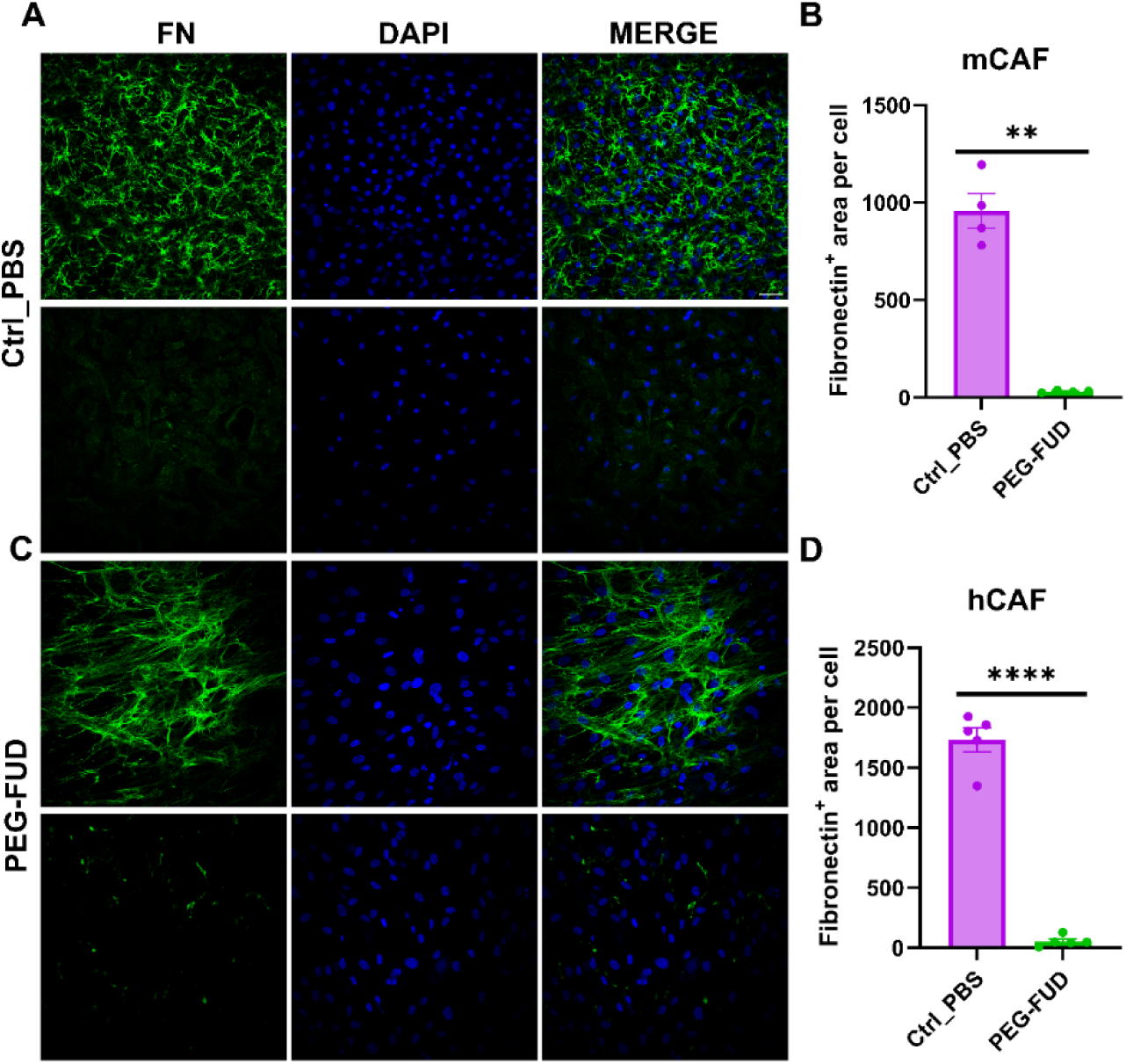
Fibronectin (FN) assembly inhibition by PEG-FUD in mCAF & hCAF. (A) Immunofluorescence staining of FN assembled by mCAF 48 hr after incubation of a vehicle control (Ctrl_PBS) or PEG-FUD (500 nM each). Scale bar = 100 μm. (B) FN (+) area changes measured by fluorescence-positive area per number of cells (nuclei). **P<0.005, n=4 per group. (C) Immunofluorescence staining of FN assembled by hCAF 48 hr after incubation of a vehicle control (Ctrl_PBS) or PEG-FUD (250 nM each). (D) FN (+) area changes measured by fluorescence-positive area per number of cells (nuclei). ****P<0.0001, n=5 per group.

### FN is highly expressed in 4T1 mammary tumors compared to other organs

FN expression is significantly increased in human breast tumors compared to normal breast tissue and is correlated with poor prognosis(43,44). To investigate the therapeutic potential of PEG-FUD to inhibit FN assembly, we chose the aggressive 4T1 pre-clinical murine model that mimics human TNBC(45). Before testing the therapeutic efficacy of PEG-FUD, we set out to validate and quantify the increase in circulating and peritumoral FN in the 4T1 tumor model. To accomplish this goal, we quantified FN levels in plasma and major organs of normal (non-tumor bearing) and tumor-bearing mice. Plasma FN (pFN) levels were similar in normal mice and in 4T1 tumor-bearing mice (Fig 2A). However, when assessing FN in major organ tissues, we found both soluble FN (sFN) and assembled / insoluble FN (iFN) levels were significantly increased in tumor tissues compared to all other major organs in both normal and tumor-bearing mice (Fig 2B-C). Specifically, 4T1 mammary tumors had more than 20-fold higher FN than normal mammary glands. Analysis of FN in the kidney, heart, liver, and spleen tissue revealed no significant difference in sFN or iFN between tumor-bearing and non-tumor-bearing mice. Notably, the lung tissue of 4T1 tumor-bearing mice had a trend toward increased iFN compared to the lungs of non-tumor-bearing mice. Similarly, others have identified an increased accumulation of total FN in metastatic 4T1 lesions isolated from multiple organ sites of tumor-bearing mice compared to non-tumor-bearing normal organs(46). Overall, our results validate that 4T1 tumors are high in sFN and iFN expression, making it a suitable model to study the anti-cancer effects of inhibiting FN assembly with PEG-FUD.

**Figure 2:**
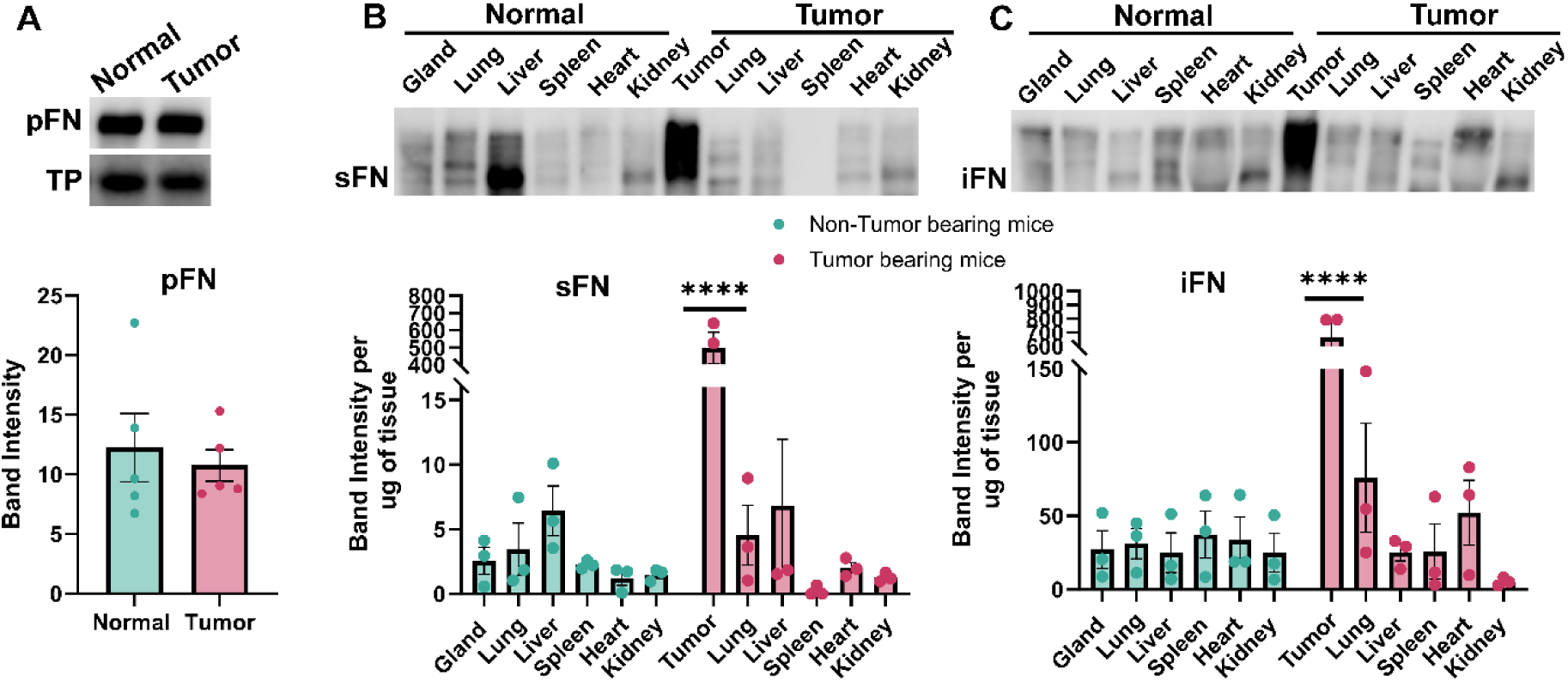
High FN expression in 4T1 mammary tumors compared to other organs. 9-week-old female BALB/c mice were injected with 2×10^5^ 4T1 cells into the fourth mammary fat pad and allowed to grow for 4 weeks. (A) Representative western blot and quantification of plasma FN (pFN) in normal, non-tumor bearing mice and in 4T1 tumor bearing mice. A representative total protein (TP) band is shown as a loading control. n=5 per group (B-C) Representative western blot and quantification of soluble FN (sFN) and insoluble FN (iFN) in multiple tissues of normal and 4T1 tumor bearing mice. The term "gland and tumor" refers to the mammary gland and mammary tumor. ****P<0.0001, n=3 per group

### PEG-FUD treatment reduced 4T1 tumor growth and decreased intratumoral FN levels

Previously we have demonstrated that PEG-FUD targets and accumulates in 4T1 mammary tumors compared to other organs using multiple imaging modalities(36). To investigate the therapeutic efficacy of targeting FN in the TME, we treated 4T1 orthotopic mammary tumors with the FN- targeting peptide, PEG-FUD, as an anti-cancer agent. One-week after 4T1 cancer cell injection into the right 4^th^ mammary fat pad, when the average tumor size reached ∼50 mm^3^, mice were subcutaneously injected with either vehicle control (Ctrl_PBS) or 12.5 mg/kg (in peptide equivalent) of PEG-FUD, PEG-mFUD or PEG-FmFUD every 48hr for a total of 10 doses (Fig 3A). The mFUD and FmFUD control peptides are mutated versions of WT FUD with 7 or 10 mutation sites, respectively (Supplemental Fig 1A). Our results showed that PEG-FUD treatment slowed tumor growth after just 4 doses compared to the control groups. Similarly, at endpoint (10 doses) PEG-FUD significantly reduced tumor volume compared to Ctrl_PBS and PEG-FmFUD (Fig 3B-C). The control peptide, PEG-mFUD, had a partial effect in reducing tumor growth, while tumor growth in the PEG-FmFUD treated mice was similar to the vehicle, Ctrl_PBS-treated mice. This was not surprising, as we previously showed that PEG-mFUD partially inhibits b-FUD binding to FN at high concentrations *in vitro*(36), and the partial binding of PEG-mFUD to FN might be explained by the remaining (non-mutated) binding sites for FN present in PEG-mFUD (Supplemental Fig 1A). This is further supported by the result from the fully mutated control peptide, PEG-FmFUD, which has additional mutations in FN binding sites and did not bind to FN in enzyme-linked competitive binding assay, suggesting that the additional mutations reduced binding to FN *in vitro* (Supplemental Fig 2A).

**Figure 3:**
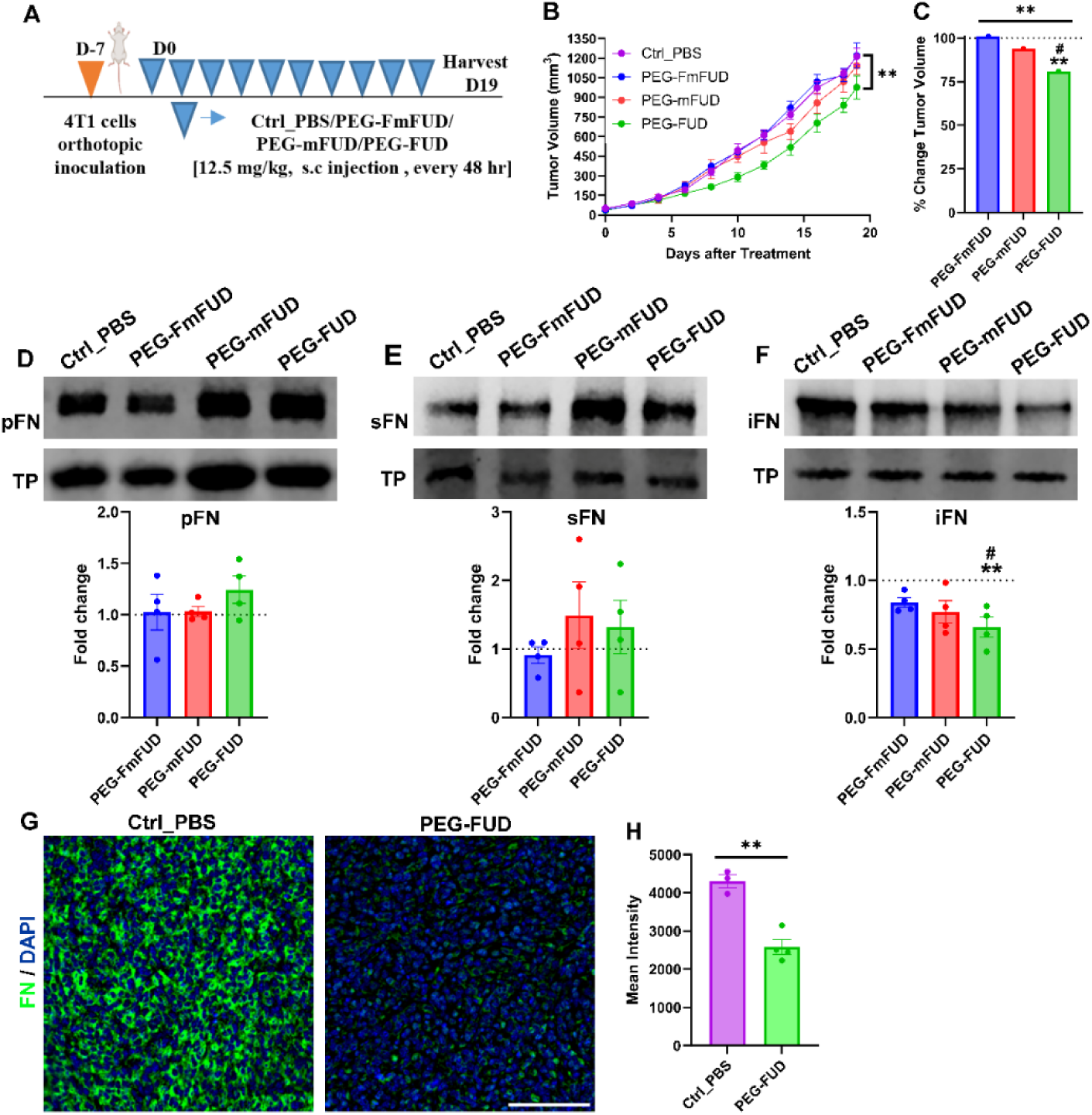
PEG-FUD significantly reduced 4T1 tumor growth and FN in the TME. (A) Experimental design for therapeutic treatment. 9-week-old, female BALB/c mice were injected with 2×10^5^ 4T1 cells into the 4th mammary fat pad. Mice were subcutaneously (s.c) treated with vehicle control (Ctrl_PBS), PEG-FmFUD, PEG-mFUD, or PEG-FUD (10 doses of 12.5 mg/kg peptide) when tumors reached ∼50 mm^3^. (B) 4T1 tumor growth post Ctrl_PBS & peptide treatments. **P<0.005, n=4 per group. (C). % Change in 4T1 tumor volume from Ctrl_PBS to mice treated with PEG-FmFUD, PEG-mFUD, or PEG-FUD. # indicates significant fold change relative to significance from vehicle control (Ctrl_PBS). **P<0.005, n=4 per group. (D-F) Representative western blots and quantification of pFN, sFN, and iFN post Ctrl_PBS and peptide treatments. A representative total protein (TP) band is shown as a loading control. The graphs represent normalized band intensities to Ctrl_PBS treatment. # indicates significance from vehicle control (Ctrl_PBS) **P<0.005, n=4 per group (G-H) Immunofluorescence staining and quantification of FN in 4T1 tumors from Ctrl_PBS and PEG- FUD treated mice (6 doses of 12.5 mg/kg peptide). Scale bar = 100 μm. **P<0.005, n=3-4 per group.

As PEG-FUD binds to the 70-kDa region of cellular and plasma FN, we evaluated the pharmacological effect of PEG-FUD on both plasma and intratumoral FN. First, we evaluated circulating FN in the plasma isolated from whole blood using western blot analysis. Results showed no changes in pFN levels between Ctrl_PBS and peptide treatment groups (Fig 3D). To detect FN in tumors, flash frozen tumor tissue was lysed into two separate fractions, DOC-soluble fraction containing sFN and DOC-insoluble fraction containing assembled iFN(47). Again, we quantified FN levels in each tumor lysate fraction by western blot. As expected, 10 doses of PEG- FUD significantly reduced iFN while no changes in sFN were observed (Fig 3E-F). As mentioned previously, the shift in tumor growth rate in response to PEG-FUD occurred early in the treatment time course. Therefore, we evaluated a subset of tumors from the Ctrl_PBS and PEG-FUD groups at an earlier treatment time point (after 6 doses). Total FN levels in these tumors were assessed by immunofluorescence staining. After just 6 doses of PEG-FUD, image analysis demonstrated a significant decrease in the levels of total FN as quantified by mean fluorescent intensity per field of view (Fig 3G-H).

FN is known to function as a provisional matrix for the deposition and structural organization of additional fibrotic matrix proteins(24). Thus, the reduction in iFN levels in PEG-FUD treated tumors led us to investigate potential changes in other peritumoral ECM proteins. Furthermore, previous *in vivo* studies using FUD as an anti-fibrotic agent in other fibrotic diseases demonstrated that FUD not only blocked FN assembly, but the treatment was accompanied by a reduction in collagen deposition(31,34). To assess alterations in other ECM proteins in the 4T1 TME post-PEG-FUD treatment, we probed tumor lysates for FN-interacting matrix proteins that we and others have identified in the 4T1 tumor model(48–50). We investigated the levels of collagen (COL1A1), tenascin-C (TNC) and periostin (POSTN) in the DOC-insoluble fraction of tumor lysates by western blot. Surprisingly, our findings showed no significant changes in COL1A1, TNC or POSTN (Supplemental Fig 3A-C). While the protein levels of COL1A1 remained unchanged with PEG-FUD treatment, the possibility remained that collagen fiber architecture may be altered due to the reduction in assembled FN. To evaluate collagen fiber architecture, we stained sections with PSR and conducted second harmonic generation imaging of collagen on thick tumor sections. Interestingly, PEG-FUD treatment did not alter collagen fiber area, nor did we observe any apparent difference in fiber architecture in any of the treatment groups (Supplemental Fig 3D- E). One possibility for this result may be due to the reduced, but not eliminated, level of iFN, which allows for the residual FN fibrils to act as a template for collagen and other ECM deposition in the TME.

### No significant toxicities were detected in PEG-FUD treated mice

Drug-induced toxicity is a major concern in developing new treatments for breast cancer patients. To validate the safe use of PEG-FUD as a therapeutic agent, we conducted a series of standard toxicity and histologic assessments. No significant body weight changes or signs of distress were observed in mice within Ctrl_PBS or peptide treatment groups (Fig 4A). The spleen weights of control and peptide treated mice were not significantly different (Fig 4B). CBC analysis indicated no changes in white blood cells (WBC), lymphocytes (LYM), monocytes (MON), or neutrophils (NEU) (Fig 4C). Additionally, platelet levels were noted to be higher with PEG-FUD treatment although not statistically significant from control groups (Fig 4D). The platelet count after PEG- FUD treatment remained within the typical range of platelet counts reported in other studies utilizing BALB/c mice(51,52). Histopathologic analysis of kidneys identified mild to moderate intracytoplasmic vacuolation within cortical tubular epithelial cells in all peptide treatment groups. This vacuolar change was not observed in the vehicle treated, Ctrl_PBS group (Supplemental Fig 4A). Prior reports show that PEG-linked proteins can induce renal tubular vacuolation at high dose or repetitive low doses, yet modifications were not linked to any changes in clinical pathology or functional indicators(53). Histopathologic analysis of liver showed mild extramedullary hematopoiesis (EMH) in both Ctrl_PBS and peptide treatment groups, with no noted difference in abundance between groups, suggesting the findings may be due to the 4T1 tumor model rather than treatment (Supplemental Fig 4B).

**Figure 4:**
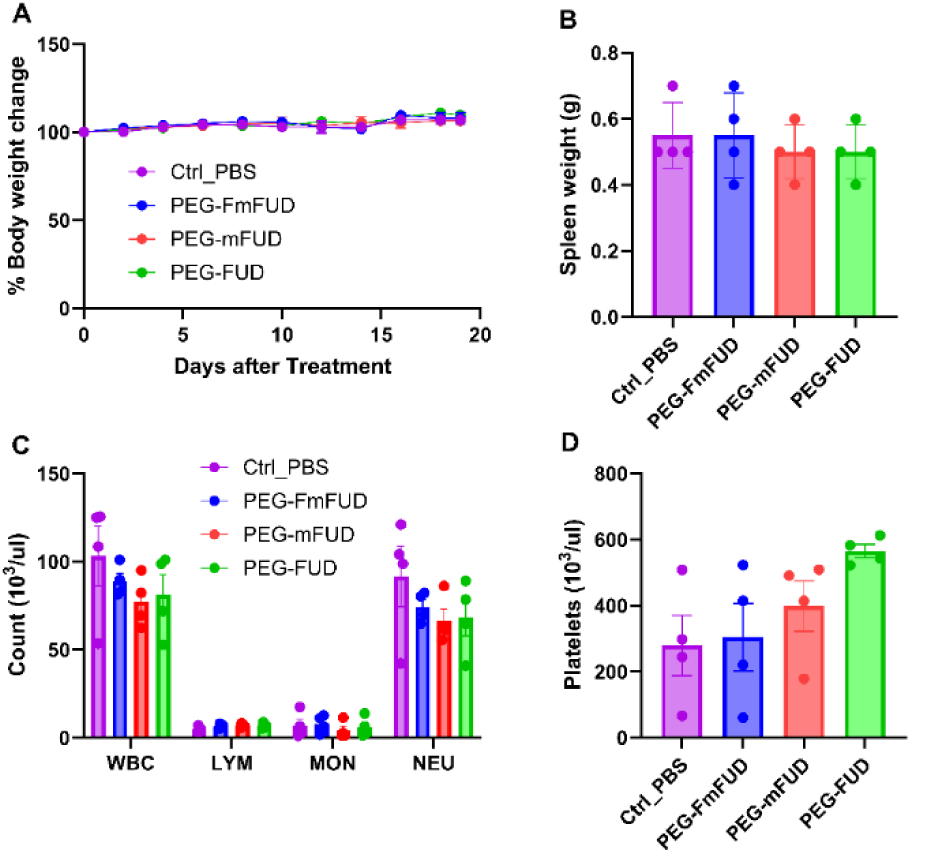
No changes in toxicity profile post PEG-FUD treatment. 2×10^5^ 4T1 tumor cells were injected into the 4th mammary fat pad and allowed to grow to 50 mm^3^. Then animals were administered vehicle control (Ctrl_PBS) or 10 peptide treatments via subcutaneous injection every 48 hr. (A) Time course of body weight measurements collected at every treatment period. (B) Quantification of spleen weight measurements at the end of treatment. (C) Complete blood count analysis of white blood cells (WBC), lymphocytes (LYM), monocytes (MON), or neutrophils (NEU) at therapeutic end point. (D) Platelet levels post Ctrl_PBS and peptide treatments. n=4 per group

### PEG-FUD treatment did not affect cell survival or proliferation signaling

FN is known to initiate signaling through integrin adhesion to activate AKT-mediated cell survival pathways and cell proliferation via the MAPK/ERK pathway. Changes in one or both of these pathways could play a role in the observed reduction in tumor size in PEG-FUD-treated mice. To examine the effect of blocking FN assembly by PEG-FUD, we quantified cell survival signaling by assessing total and phosphorylated AKT (p-AKT) levels in tumor lysates. Surprisingly, no changes were observed in total AKT or activated p-AKT (Fig 5A). Next, we assessed ERK-regulated cell proliferation by quantifying total and phosphorylated ERK (p-ERK). Similar to our results with AKT activity, we found no changes in ERK activation in response to PEG-FUD treatment (Fig 5B). Additionally, we examined two well-known cell proliferation markers, PCNA in tumor lysates after 10 doses of peptide treatment and the area of Ki67+ cells after 6 dose of peptide treatment, using western blotting and immunofluorescence, respectively. Consistent with the above findings, we did not detect a change in either marker of cell proliferation in response to PEG-FUD treatment compared to control groups (Fig 5C-E). Although these findings contradict previous research showing that inhibiting FN decreased proliferation(28,30,31), this may be explained by a difference in orthotopic tumor models or differences in growth factors present in the TME of 4T1 tumors that could maintain proliferation signaling.

**Figure 5:**
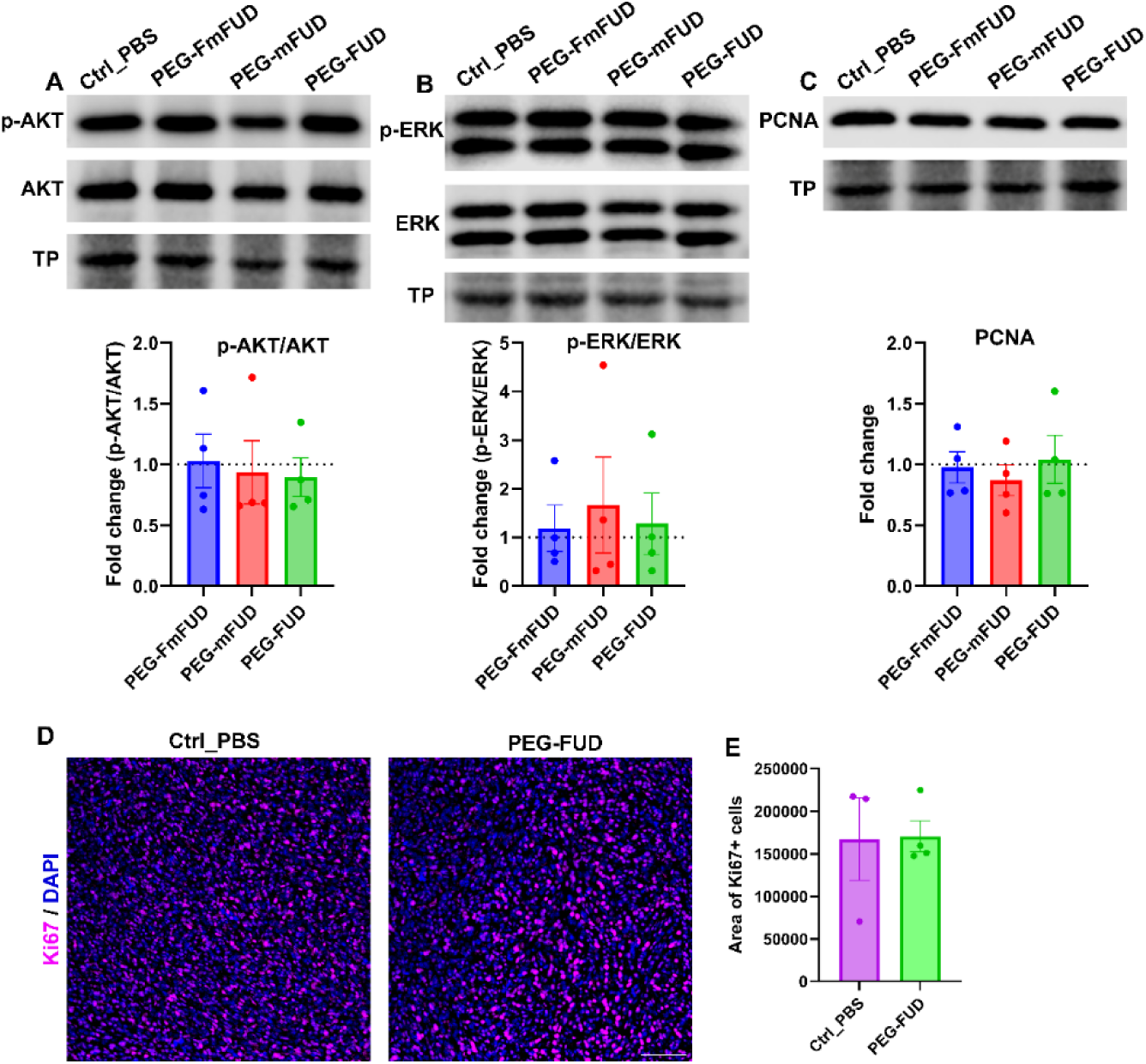
No changes in cell survival / proliferation signaling post PEG-FUD treatment. (A-C) Representative western blots and quantification of proliferation markers AKT, ERK and PCNA post vehicle control (Ctrl_PBS) and peptide treatments (10 doses of 12.5 mg/kg peptide). A representative total protein (TP) band is shown as a loading control. The graphs represent normalized band intensities to Ctrl_PBS treatment. (D) Representative immunofluorescent images stained for proliferation marker Ki67 post Ctrl_PBS and PEG- FUD treatment (6 doses of 12.5 mg/kg peptide). (E) Quantification of Ki67+ area from D.

### Treatment with PEG-FUD reduced α5 integrin and FAK protein expression and promoted caspase-3 mediated cell death

Next, we sought to interrogate FN-mediated focal adhesions and downstream apoptotic signaling as part of the mechanism by which PEG-FUD therapy inhibits tumor growth. Integrin α5β1 is the primary receptor for FN that binds sFN and initiates the assembly of FN into a fibrillar matrix(54). In addition, the binding of integrins to FN activates intracellular signaling pathways downstream of focal adhesions(55). To determine whether adhesion-mediated signaling is disrupted in response to PEG-FUD, we quantified the level of α5 and β1 integrins by western blot analysis. PEG-FUD treatment resulted in a significant reduction in the tumor levels of integrin α5 (Fig 6A). In contrast, PEG-FUD therapy did not affect β1 integrin (Fig 6B). β1 integrins form complexes with multiple alpha integrins subunits to bind several ECM proteins found in the TME(56). Thus, the lack of β1 integrin specificity for FN is one possible rationale for the modulation of α5, but not β1 integrin, in response to PEG-FUD treatment. Due to the reduction in iFN and α5 integrin levels, we next evaluated the effect of PEG-FUD on the focal adhesion signaling through focal adhesion kinase (FAK). FAK is a master regulator of cell adhesion signaling upon integrin activation and clustering(57). Strikingly, we observed a significant reduction in total FAK protein levels in PEG- FUD treated tumors compared to controls (Fig 6C). Taken together, these findings indicate that inhibiting FN assembly with PEG-FUD alters FAK-dependent integrin-mediated cell adhesion.

**Figure 6:**
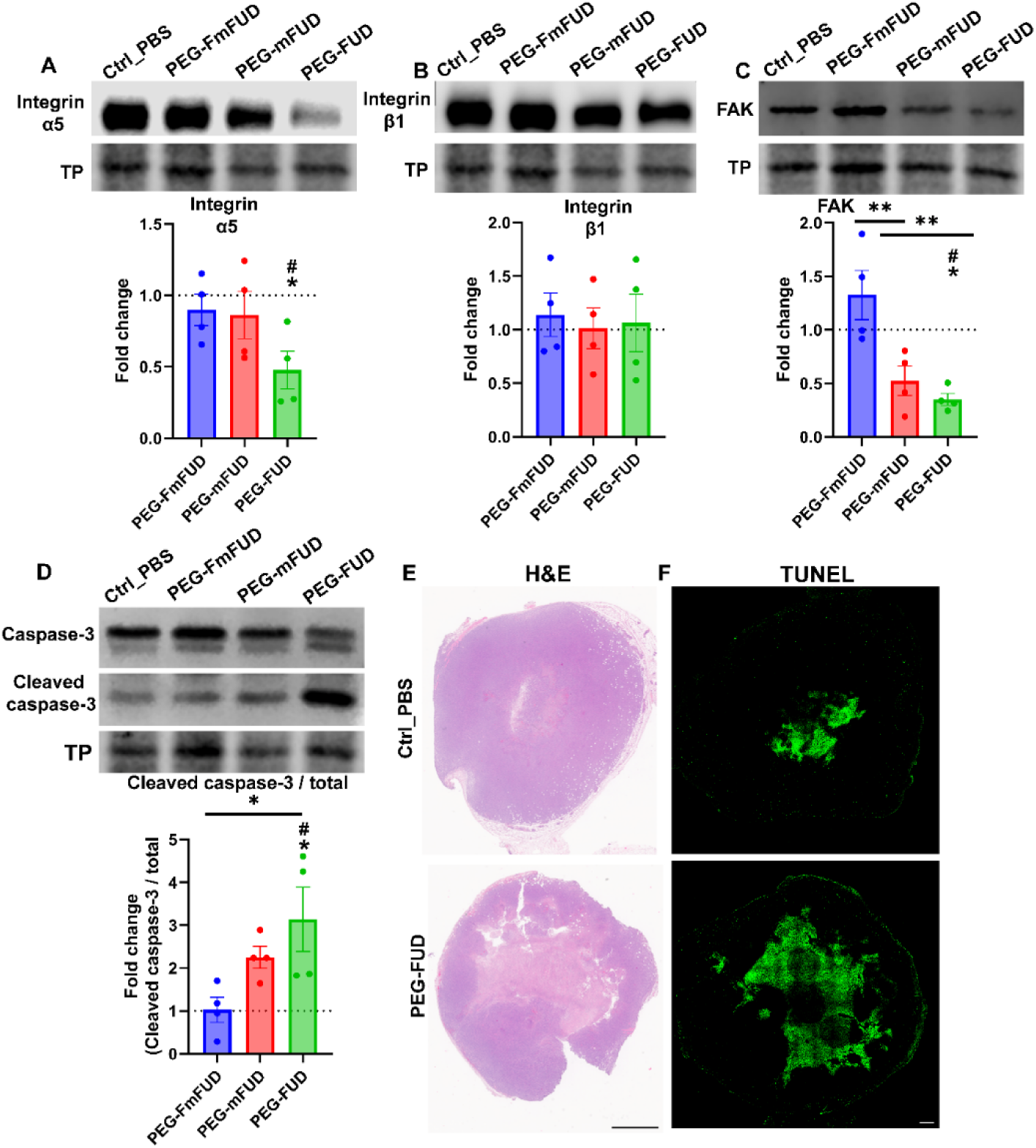
PEG-FUD alters integrin mediated adhesion signaling and cell death. (A-D) Representative western blots and quantification of Integrin α5, Integrin β1, FAK, caspase-3 and cleaved caspase-3 from 4T1 tumor lysates treated with vehicle control (Ctrl_PBS) and peptides (10 doses of 12.5 mg/kg peptide). A representative total protein (TP) band is shown as a loading control. The graphs represent normalized band intensities to Ctrl_PBS treatment. # indicates significant fold change relative to significance from vehicle control (Ctrl_PBS). *P<0.05, **P<0.005, n=4 per group (E) Representative images of H&E staining and TUNEL staining of 4T1 tumor sections from mice treated with Ctrl_PBS and PEG-FUD (6 doses of 12.5 mg/kg peptide). H&E Scale bar = 100 μm; TUNEL Scale bar = 500 μm.

In the literature, FN knockdown, blockade of FN receptor integrin α5β1, and alterations in FAK have been shown to induce anoikis, a type of programmed cell death that results from a loss of cell-ECM attachment(58,59). To investigate the effect of FN inhibition by PEG-FUD in mediating cell death, we probed for cell death markers at both early and late treatment time points (6 or 10 doses of PEG-FUD). We confirmed the induction of cell death by PEG-FUD by a statistically significant increase in cleaved caspase-3 levels from whole tumor lysates collected from mice that received 10 doses of PEG-FUD compared to control-treated tumor groups (Fig 6D). Tumors harvested from mice that received 6 doses of PEG-FUD showed a qualitative increase in the presence of tumor necrosis identified in H&E stained sections (Fig 6E). Additionally, representative serial sections from each treatment group were assessed by TUNEL assay. We identified large TUNEL positive regions at the core of PEG-FUD treated tumors that are not observed in the vehicle control (Ctrl_PBS) treated tumors (Fig 6F). Altogether, these results suggest that PEG-FUD induced cell death by inhibiting FN-integrin-mediated cell adhesion.

### PEG-FUD primes the mammary TME for enhanced anti-tumor efficacy of doxorubicin

The identification of PEG-FUD-mediated tumor cell death led us to hypothesize that PEG-FUD not only has intrinsic anti-cancer potential but may also synergistically improve the tumor response to chemotherapy. Others have demonstrated that the overexpression of FAK, which is identified in multiple tumors, plays a critical role in suppressing tumor cell death and facilitating resistance to Dox(60,61). Based on these findings and our evidence for PEG-FUD to robustly inhibit FAK protein levels in the 4T1 TME, we tested the ability of PEG-FUD pre-treatment to improve the response of a commonly used chemotherapeutic drug, Dox, in the 4T1 tumor model. 4T1 orthotropic tumors were established similarly to the PEG-FUD monotherapy approach with treatment starting when the average tumor size reached ∼50 mm^3^. Mice were randomized into treatment groups and subcutaneously injected with either vehicle control (Ctrl_PBS) or 12.5 mg/kg (in peptide equivalent) of PEG-FUD or PEG-FmFUD every 48 hr for 4 doses. At the 4-dose timepoint (when a shift in tumor growth was noted, Fig 3B), we initiated intraperitoneal administration of Dox at 5mg/kg every 72 hr with continued peptide treatment (experimental design outlined in Fig 7A). Combinatorial treatment of PEG-FUD and Dox resulted in a significant reduction in tumor growth by ∼60% to vehicle control and by an additional ∼30% to Dox alone, demonstrating that PEG-FUD co-treatment increased the efficacy of Dox therapy in the 4T1 TNBC model (Fig 7B-C). Mice treated with Dox exhibited loss of body and spleen weight due to the well-known systemic toxicity of Dox, but no additive effect was detected with co-treatment with PEG- FUD (Fig 7D-E). These findings provide evidence that reducing FN with PEG-FUD improves chemotherapeutic response.

**Figure 7:**
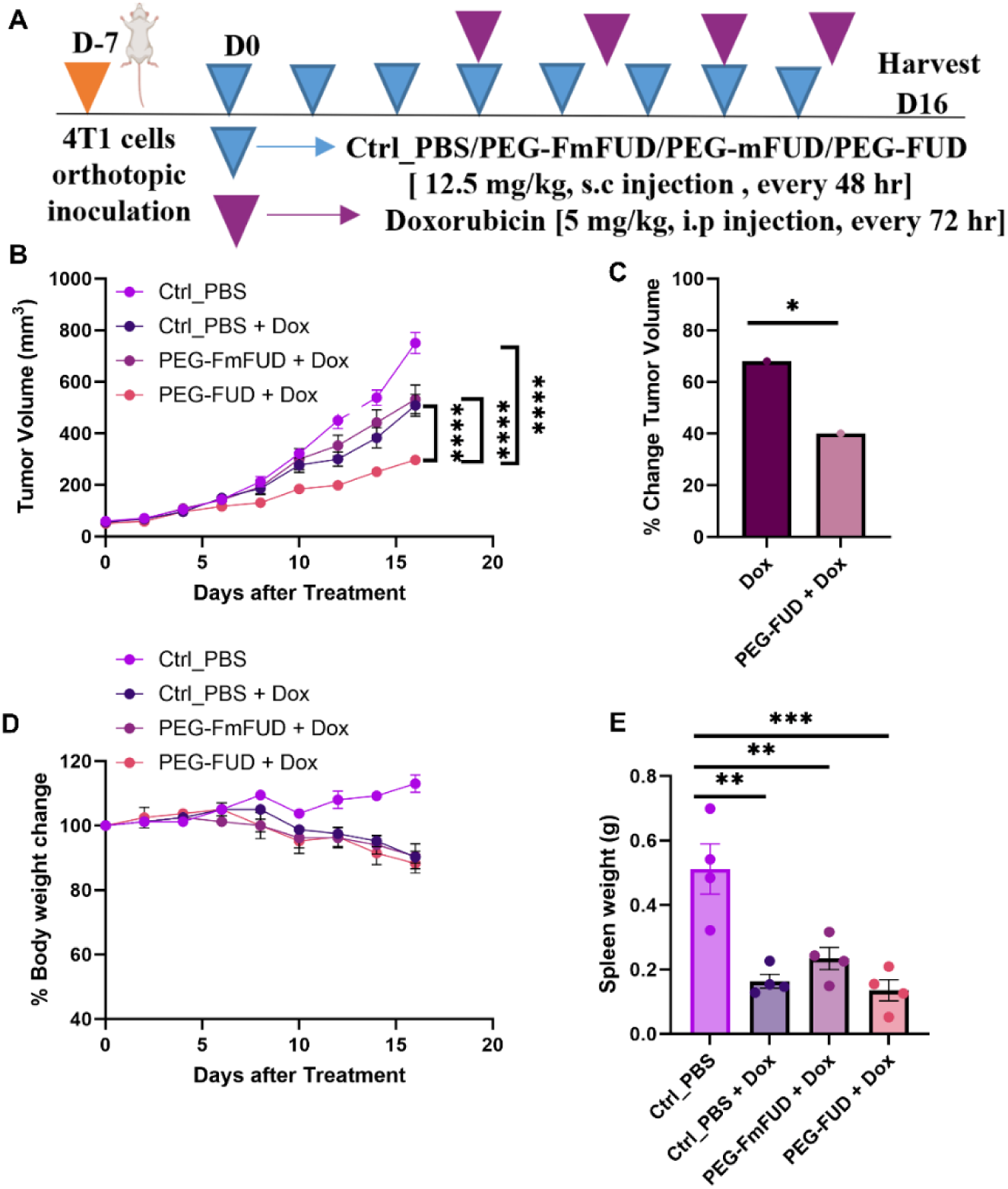
PEG-FUD combined with Doxorubicin resulted in enhanced efficacy without additional toxicity. (A) Experimental design for therapeutic treatment. 9-week-old, female BALB/c mice were injected with 2×10^5^ 4T1 cells into the 4th mammary fat pad. Mice were subcutaneously (s.c) treated with 4 doses of vehicle control (Ctrl_PBS), PEG-FmFUD, or PEG-FUD at 12.5 mg/kg every 48 hrs once tumors reached ∼ 50 mm^3^. Upon the 4^th^ dose of PEG-FUD, Doxorubicin (Dox) was administered intraperitoneally (i.p) at 5mg/kg every 72 hr with continued peptide treatment until day 16 post 4T1 tumor cell inoculation. (B) 4T1 tumor growth post Ctrl_PBS and combined treatments. ****P<0.0001, n=4 per group. (C). % Change in 4T1 tumor volume from Ctrl_PBS to mice treated with Dox and PEG-FUD + Dox. *P<0.05, n=4 per group. (D) Time course of body weight measurements collected at every treatment period post vehicle control (Ctrl_PBS), Dox and peptide + Dox combination treatments. (E) Quantification of spleen weights post Ctrl_PBS, Dox and peptide combination treatments at the end of treatment period. **P<0.005, ***P<0.001, n=4 mice per group.

## Discussion

In this study, we investigated the anti-cancer effects of targeting FN matrix assembly in the TME using PEG-FUD. Several FN-targeting strategies have been developed and demonstrated promise in the diagnosis, therapy, and image-guided interventions of a variety of malignancies(62). For example, antibodies that detect the EDA domain of cellular FN have shown promise for solid tumor diagnostics(63). Additional studies expanded on the utility of targeting FN by developing antibody-drug conjugates using an EDA-antibody with anti-IL-2(64), EDA expressing CAR-T cells(65), or developing fibrin-FN binding peptides conjugated with Dox(66) for enhanced therapeutic efficacy and reduced systemic toxicities. Some of these approaches made it to early-phase human trials, however, they have yet to be FDA-approved for the clinical treatment of breast cancer. Thus, developing new therapeutic approaches to target FN in the TME is both timely and promising.

Here, we take advantage of targeting FN with PEG-FUD. PEG-FUD binds with high affinity to the 70-kDa N-terminus and inhibits the assembly of all sources of FN (cellular and plasma derived FN). This is essential since both sources of FN can be deposited in the tumor matrix(67,68). We previously reported that PEG-FUD accumulates and is maintained in 4T1 tumors for prolonged periods compared to FUD alone(36). Our current *in vitro* data demonstrate that PEG-FUD successfully inhibited the assembly of endogenous FN assembled by mammary CAFs (Fig 1). In addition, we demonstrated that PEG-FUD is effective when given in therapeutic doses to reduce the assembly of tumor FN (decreased iFN levels) and to suppress 4T1 tumor growth (Fig 3). Therefore, like prior studies we confirmed that the PEGylation of the FUD peptide retained the ability to block FN assembly(33,34,36). Together these data demonstrate the utility of PEG-FUD not only for tumor diagnostics but also as an anti-breast cancer therapeutic.

Mechanistically we determined that PEG-FUD treatment significantly reduced levels of α5 integrin, a major adhesion receptor for FN. Integrin binding to FN mediates intracellular signaling to regulate cellular processes such as cell growth, proliferation, and migration(69–71). Despite the significant reduction in α5 integrin and FAK, we did not observe differences in proliferation or survival signaling through ERK or AKT, respectively. This result contradicts previous observations that FUD/puR4 peptide decreased ERK phosphorylation without affecting FAK or AKT phosphorylation(31). However, the difference between our findings and those of Ghura et al. may be explained by differences in model systems (immune competent vs immune compromised), treatment conditions (PEG-FUD vs FUD), and identification of altered signaling pathways (within tumors vs in cell culture). Furthermore, ERK and AKT activation can occur independent of FN- FAK mediated signaling. In support of this hypothesis, a study utilizing an integrin antagonist 1a- RGD in glioblastoma cells showed that by inhibiting cell adhesion, FAK activation decreased without altering ERK and AKT-dependent pathways(72). It is also plausible that the reduction in iFN could release growth hormones normally sequestered in the matrix that become available to bind and activate pathways downstream of growth factor receptors(73–76).

In our study, the loss of α5 and FAK in the TME in response to PEG-FUD is notable. While we do not know the temporal sequence of α5 integrin vs FAK down-regulation, the functional inhibition of α5β1 integrin activation has been shown to reduce total FAK expression and protein levels(77). Others have also demonstrated that caspase-mediated FAK cleavage (degradation of FAK) causes focal adhesion complexes to disassemble(78), suggesting the activation of caspase signaling could reduce levels of FAK and α5 integrin. Importantly, the disassembly of focal adhesions down stream of caspase activation interferes with integrin-mediated survival signaling, thereby propagating the cell death program(78). In line with these studies, we identified an increase in TUNEL-positive staining and caspase-3 mediated cell death in PEG-FUD-treated tumors. Multiple studies have shown that integrin adhesion to FN can induce anoikis resistance and reduce apoptosis(58,79). Moreover, FAK itself has been identified to play a critical role in suppressing cell death(80–83). FAK has been shown to directly bind to the death domain of receptor-interacting protein (RIP) to suppress apoptosis(61). RIP provides proapoptotic signals that are suppressed upon binding to FAK, thus blocking the FAK-RIP interaction or reducing total FAK levels shifts cells toward caspase-mediated cell death. Taken together, our findings suggest that PEG-FUD effectively blocked FN assembly within the TME resulting in decreased α5 integrin and FAK levels, leading to apoptosis via the activation of caspase-3 cleavage. This results in an overall reduction in tumor growth, demonstrating PEG-FUD as an effective single agent therapy to target pre-clinical breast cancer.

Our characterization of the tumor response to PEG-FUD also demonstrates the promise of targeting FN to improve standard-of-care cancer therapies. For many solid tumors, including TNBC, the first-line treatment is chemotherapy. However, the ECM of solid breast tumors activates mechanisms of drug resistance that significantly impede clinical therapies(84–86). Other pre-clinical mammary tumor studies have shown that Pirfenidone, an FDA approved antifibrotic drug for idiopathic pulmonary fibrosis, normalizes the tumor ECM for enhanced chemotherapy response, demonstrating the value of ECM normalization(87). This study focused on the action of pirfenidone to block TGF-b signaling and downstream ECM deposition, which led to increased drug penetration. Our data suggest that the anti-fibrotic impact of PEG-FUD not only reduces iFN within tumors but also alleviates ECM-mediated mechanisms of drug resistance through the reduction of FAK. In line with our findings, van Nimwegen et al. illuminated a mechanism by which adhesion-mediated stress fiber formation and FAK activation suppressed Dox-induced apoptosis(60). Based on these results, we hypothesized that by reducing tumoral FN and inhibiting α5 integrin and FAK, PEG-FUD sufficiently primed the TME for improved chemotherapy response. Indeed, when 4T1 tumors were primed with PEG-FUD followed by combined treatment with Dox + PEG-FUD we quantified a 60% decrease in tumor growth compared to vehicle control and 30% compared to Dox monotherapy. Notably, combinatorial treatment of PEG-FUD and Dox did not increase the systemic toxicity of Dox (Fig 7). Thus, PEG-FUD therapy is an effective strategy to improve the efficacy of standard-of-care chemotherapy in 4T1 TNBC model.

## Conclusion

In summary, we have demonstrated the efficacy of PEG-FUD to target FN in 4T1 triple-negative tumors and improve response to standard chemotherapy. We have characterized changes in the TME in response to PEG-FUD treatment and identified a mechanism where the reduction in iFN leads to a significant decrease in α5 integrin and FAK levels. Further, these changes in cell adhesion increased tumor cell death and blocked tumor growth. Strikingly, the tumor response to PEG-FUD therapy was sufficient to prime the TME into a non-tumor-supporting microenvironment and significantly enhance the efficacy of Dox therapy. As we previously demonstrated PEG-FUD can also target the metastatic microenvironment(36), our future studies will explore PEG-FUD as an effective treatment for metastatic breast cancer.

## Acknowledgments

This work was supported by the National Cancer Institute funding to SMP (R01CA206458, and R01CA179556) and R21CA252579 to SMP and GSK. Additionally, this research was supported by the University of Wisconsin-Carbone Cancer Center, More for Stage IV Research Award to SMP.

## Author contributions

MKG conceptualized the study, conducted experiments, analyzed and interpreted data, organized figures, wrote and edited the manuscript. HJL conducted experiments, analyzed data, and edited the manuscript. DRI assisted with all animal experiments. NG prepared PEGylated FUD, conducted experiments, analyzed and edited the manuscript. BB assisted with data analysis, interpretation and edited the manuscript. MH conducted the histopathologic assessment and edited the manuscript. GSK acquired funding, conceptualized the study, interpreted data, and edited the manuscript. SMP acquired funding, conceptualized the study, oversaw the experiments and data collection, interpreted analysis, wrote and edited the manuscript.

