## Supplemental Figures for "Inhibiting fibronectin assembly in the breast tumor microenvironment increases cell death and improves response to doxorubicin"

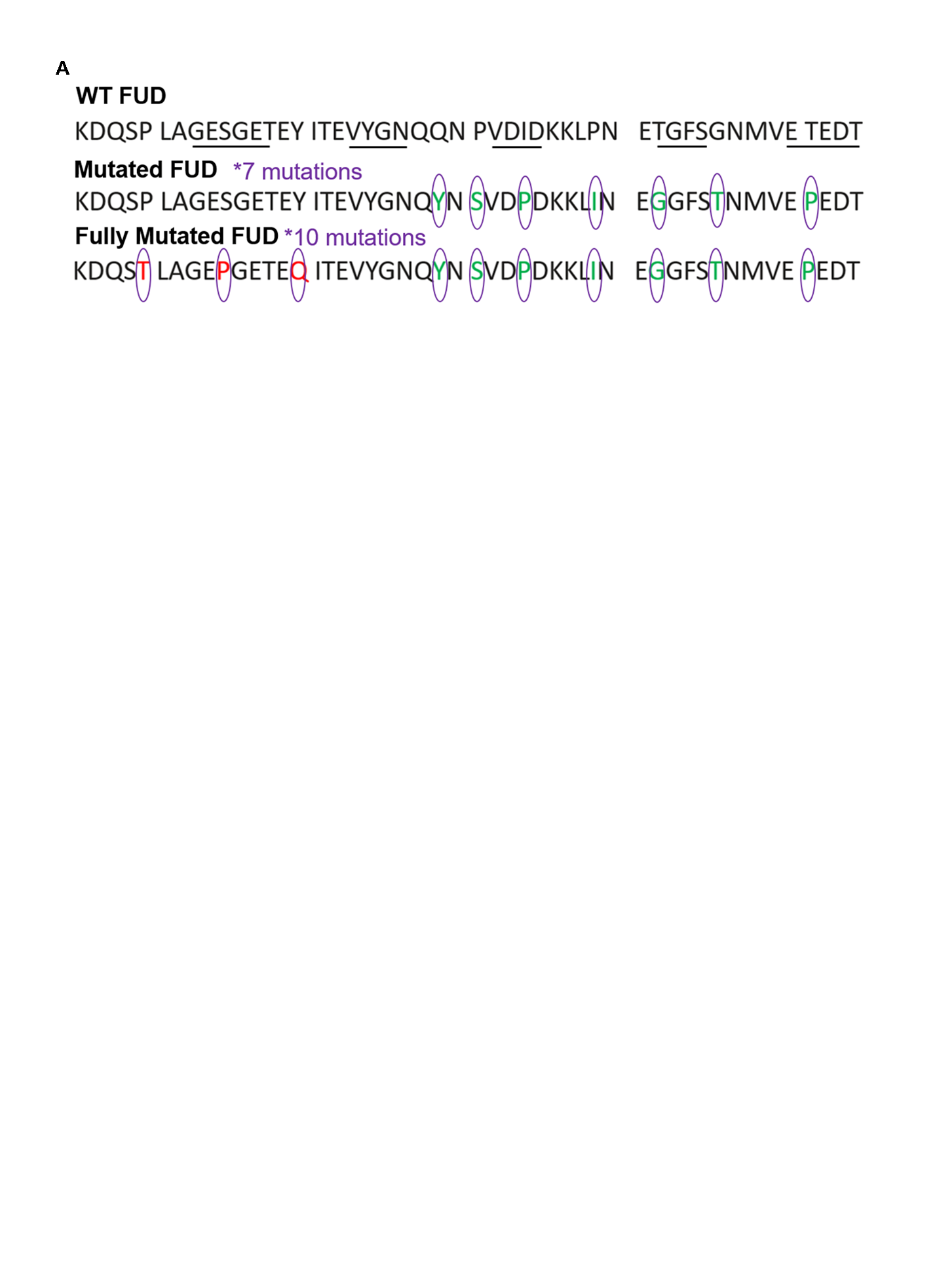

**Supplemental Figure 1.** (A) FUD and FUD derived mutated peptide sequences. Mutated amino acids of mFUD (green) and additional mutated amino acids in Fully Mutated FUD (red). Lines indicate binding sites to FN.

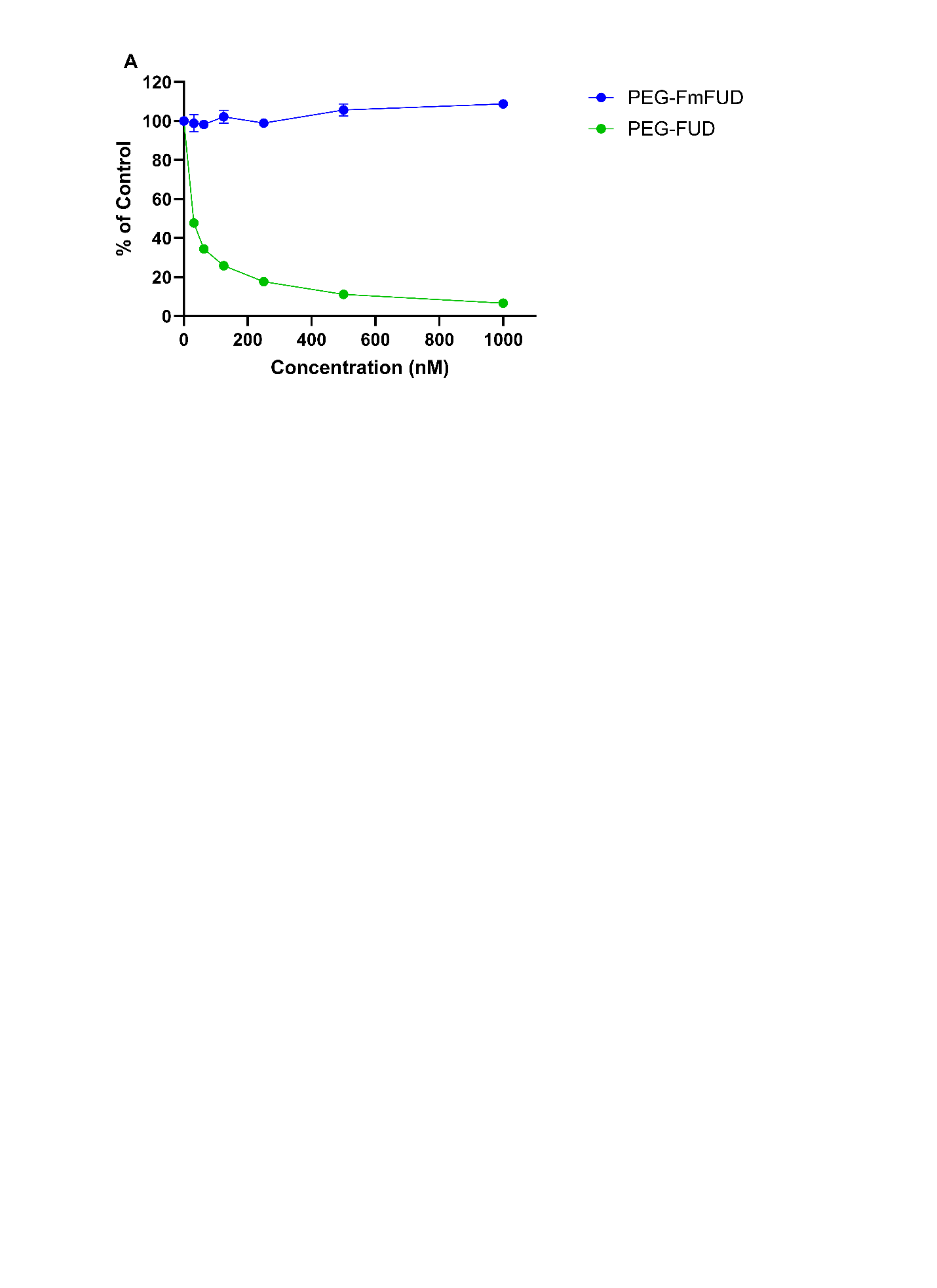

**Supplemental Figure 2.** (A) Enzyme-linked competitive binding assay showing the percent of biotinylated FUD (b-FUD) attachment to adsorbed FN when co-incubated with different concentrations of PEG-FmFUD or PEG-FUD. n= 3 per group

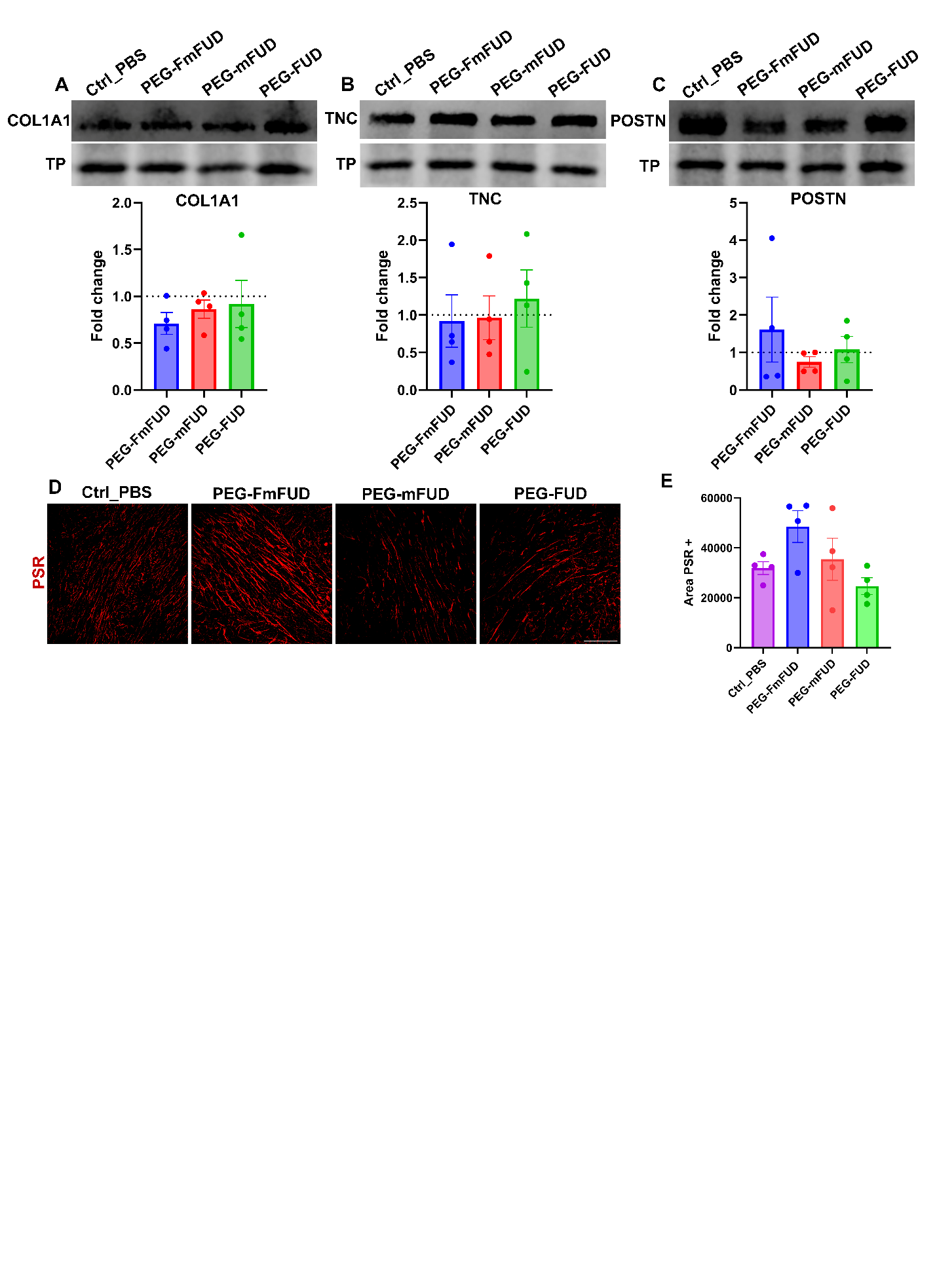
**Supplemental Figure 3.** (A-C) Representative western blots and quantification of COL1A1, TNC & POSTN from 4T1 tumor lysates treated with vehicle control (Ctrl_PBS) or peptides (10 doses of 12.5 mg/kg peptide). Representative total protein (TP) band shown as loading control. Graphs represent normalized band intensities to Ctrl_PBS treatment. (D-E) Representative picrosirius red (PSR) staining images and quantification of 4T1 tumors treated with Ctrl_PBS or peptides (10 doses of 12.5 mg/kg peptide). Scale bar = 500 μm. n=4 mice per group

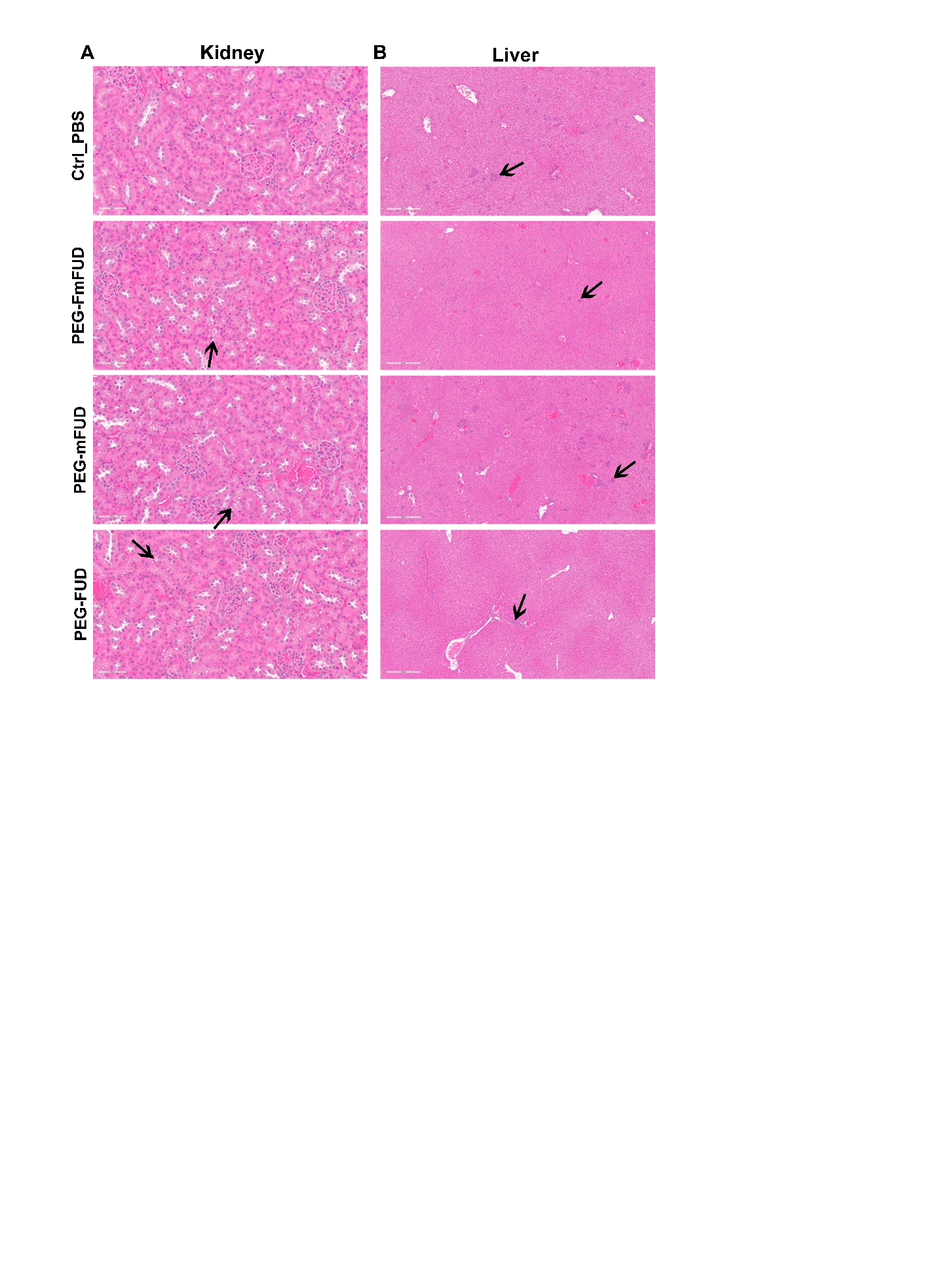

**Supplemental Figure 4.** Histopathologic evaluation of H&E stained sections of kidney and liver from mice treated with vehicle control (Ctrl_PBS) peptides (10 doses of 12.5 mg/kg peptide). (A) Representative images of kidney illustrating intracytoplasmic vacuolation of tubular epithelial cells in each treatment group (arrows), not observed in the control group. Scale bar = 60 μm. (B) Representative images of liver, illustrating multifocal extramedullary hematopoiesis (EMH) identified in each group (arrows). Scale bar = 300 μm

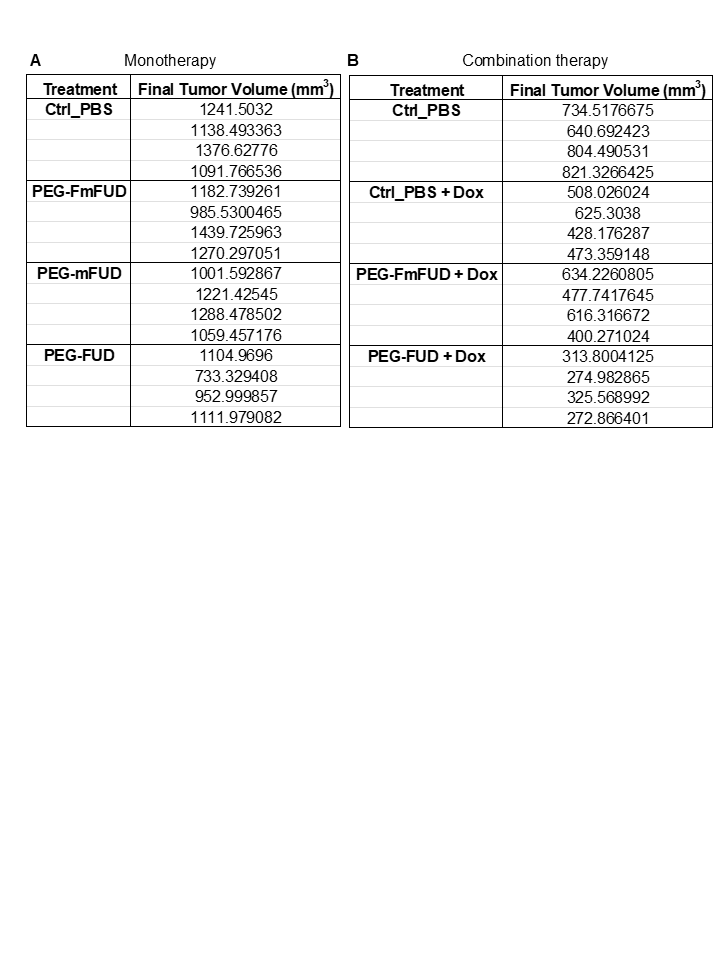

**Supplemental Table 1.** Final tumor volume measurements (A) PEG-FUD monotherapy (B) Combination therapy of PEG-FUD + Doxorubicin (Dox).
